# Specificity Profiling of the RhoGEF Domain of FP10 with Rho GTPases Involved in Cytoskeleton Remodeling from *Entamoeba histolytica*

**DOI:** 10.64898/2026.05.08.723678

**Authors:** Avinash Kumar Gautam, Preeti Umarao, Samudrala Gourinath

## Abstract

The Rho family of small GTPases plays a critical role in regulating actin cytoskeleton dynamics during endocytic processes in *E. histolytica*, including phagocytosis, pinocytosis, and trogocytosis. These proteins act as molecular switches, transitioning between inactive GDP-bound and active GTP-bound states, with guanine nucleotide exchange factors (GEFs) catalyzing this transition. Among the GEFs, EhFP10—a FYVE-domain-containing protein harbouring Dbl homology (DH) and pleckstrin homology (PH) domain was observed in phagocytosis along with seven functionally characterized Rho GTPases (EhRho1, EhRho2, EhRho4, EhRho5, EhRho6, EhRho8, and EhRho13).

To study the specificity of FP10, a combination of GEF activity, binding affinity, and molecular dynamics simulations was used to characterize the interactions between EhFP10 and seven Rho GTPases systematically. The results revealed EhRho2 as the most specific and high-affinity interactor of EhFP10, with the highest nucleotide exchange rate and lowest dissociation constant (K_D_ = 0.58 µM). Structural modeling, sequence alignment, and interaction mapping further demonstrated that EhRho2 retains critical contact residues—such as Glu33, Arg4, and Leu69—that are variably absent in other isoforms, correlating with decreased GEF responsiveness.

Molecular dynamics simulations and cross-correlation analyses supported the presence of a stable and coordinated interaction interface in the EhFP10–EhRho2 complex, distinguishing it from less active complexes. These findings indicate a highly selective GEF-GTPase module in *E. histolytica*, analogous to those in higher eukaryotes. The results uncover a potential regulatory mechanism specific to pathogenic amoebae and present EhFP10–EhRho2 as a novel therapeutic target for disrupting cytoskeleton-mediated processes crucial to virulence.

## INTRODUCTION

Small GTPase proteins are highly conserved molecular switches that play pivotal roles in cellular regulation across eukaryotes—from simple unicellular organisms like amoebae to complex multicellular organisms, including humans ^1^ and also in prokaryotes ^2^. These proteins orchestrate a wide range of cellular processes, including proliferation, differentiation, vesicle and organelle trafficking, nuclear transport, and the dynamic remodeling of the cytoskeleton ^3,4^. Their regulatory versatility lies in their ability to cycle between two distinct states: an inactive GDP-bound form and an active GTP-bound form ^5^. This cyclical transition enables them to act as binary molecular switches that control downstream signaling pathways in a temporally and spatially precise manner ^6^.

Structurally, small GTPases belong to the Ras superfamily—so named after the Rat sarcoma (Ras) oncogene, one of the first identified and studied members. The Ras superfamily is subdivided into five major subfamilies based on sequence homology and functional specificity: Ras, Rho, Rab, Arf, and Ran ^7,8^. Each subfamily is associated with a unique cellular function ^5,9,10^. For instance, Ras proteins regulate cell growth and proliferation; Rho proteins control actin cytoskeleton organization ^11,12^; Rab proteins oversee vesicle trafficking and fusion; Arf proteins are key players in vesicle formation and recycling ^13^; Ran proteins are involved in nuclear transport and mitotic spindle assembly ^14^.

All small GTPases share a highly conserved GTP-binding domain comprising five sequence motifs known as G boxes (G1–G5) ^8^. These motifs coordinate the binding and hydrolysis of guanine nucleotides (GDP and GTP) and the essential cofactor Mg^2+^, stabilizing the nucleotide-binding pocket. The Switch I region typically coordinates the purine ring of GTP. In contrast, the Switch II region undergoes significant conformational changes upon nucleotide exchange, a process central to the GTPase’s ability to transmit signals to downstream effectors ^6,7,15,16^.

Three classes of regulatory proteins tightly regulate the GTPase activation cycle: Guanine Nucleotide Exchange Factors (GEFs), GTPase-Activating Proteins (GAPs), and Guanine Nucleotide Dissociation Inhibitors (GDIs) ^17,18^. GEFs promote the exchange of GDP for GTP, thereby activating the GTPase ^19,20^. Mechanistically, GEFs interact with key structural elements of the GTPase, particularly the G2 box and switch regions, to induce conformational changes that weaken the affinity for GDP ^6,7^. The CR1 region of GEFs engages the switch regions and destabilizes GDP binding, while the CR3 region stabilizes the nucleotide-free intermediate. Once GDP is released, the high intracellular concentration of GTP facilitates its rapid binding to the now-vacant nucleotide pocket. GTP binding reorients the switch regions into an active conformation, and GEF subsequently dissociates from the complex.

Different families of GEFs exhibit specificity for distinct GTPase subfamilies. Son of Sevenless (Sos) proteins are specific for the Ras subfamily, Dbl-homology domain-containing GEFs act on Rho GTPases ^21^, Sec7 domain GEFs target Arf proteins, and EF-Tu-like GEFs facilitate nucleotide exchange in translational GTPases ^22^. Despite their diversity, all GEFS share the conserved mechanism of destabilizing GDP binding and promoting GTP loading to activate their respective GTPases ^20^.

In the protozoan parasite *Entamoeba histolytica*, small GTPases have evolved to play essential roles in modulating the actin cytoskeleton during processes such as pinocytosis, phagocytosis, and encystation, all of which are critical for virulence and survival ^23,24^. Several small GTPases and GEFs in *E. histolytica* have been functionally characterized ^25,26^. EhRho1 is involved in actin cytoskeleton remodeling and bleb formation ^27–30^. EhRho2/EhRacG^1^ localize to the phagocytic vacuole and interact with Myosin-IB, which is overexpressed during phagocytosis ^31^. EhRacA participates in receptor capping and localizes to the leading edge of the phagocytic cup, promoting F-actin polymerization and pseudopod extension. EhRho5 is dynamically activated by lysophosphatidic acid (LPA) and relocates from the cytosol to membrane structures, underscoring its involvement in vesicle trafficking and signal transduction ^32^. EhRho6 and EhRho8 are key players in actin degradation and are also essential for encystation, phagocytosis, and motility. Notably, EhRho8 is expressed 53-fold higher in cysts than in trophozoites, highlighting its role in structural reorganization during encystation ^33^. EhRho13, unlike its counterparts, is primarily involved in ribosomal protein quality control and nucleotide metabolism, negatively regulating micropinocytosis ^34^. These proteins underscore the functional diversity and complexity of small GTPases in *E. histolytica*, illustrating that a conserved molecular mechanism may be adapted for specific roles. The regulation of these GTPases via Dbl-RhoGEFs (EhGEF2, EhGEF3) is also reported to be important for the pathogen cellular processes ^35,36^.

The EhFP10 has a RhoGEF domain-containing, along with a FYVE domain, which was found to be involved in endocytosis by interacting with actin and Myosin ^37^. The FP10 contains a Dbl Homology (DH) domain and a Pleckstrin Homology (PH) domain, which together perform GEF function. These DH-PH domain-containing proteins, along with the FYVE domain, are only seen in amoebic species ^38^. Recently, we also showed that FP10 interacts with PAK4 and Rho2 (Gautam et al., 2026, accepted for publication in FEBS from our lab). But the specificity of Rho was not clear when this was observed in phagocytic cup formation, as all seven Rho GTPases were present in endocytic and phagocytic processes ^27,31–34^. Here, we studied the specificity of Rho GTPases for EhFP10 using GTPase activity, BLI interactions, complex models, and simulations to understand specificity and correlated it with residues and interactions in complex formation.

## RESULTS

All seven^i^ RhoGTPase proteins (EhRho1, EhRho2/EhRacG, EhRho4/EhRacA, EhRho5/EhRacD1, EhRho6/EhRacA2, EhRho8, EhRho13/EhRacM) were cloned into pET28b+ and purified by Ni-NTA affinity chromatography (Table S1 & Figure S1). The sequence alignment of all seven Rho GTPases (Figure 1) shows typical features of conserved G1-G5 domains and a long, flexible C-terminal region containing a CXXA motif, which is involved in post-translational modification. The C-terminal modification specifies the localization of the RhoGTPase for its function and facilitates the association of membrane lipid moieties. For nucleotide exchange, two major regions, switch-1 and switch-2, are crucial and are highly conserved in the sequence alignment. Further, the structures of six RhoGTPases were modeled (Figure S2), and Rho1 is shown in GDP-bound form (PDB:3REF) and GTP analog-bound form (PDB:3REG) from an earlier crystal structure^39^ (Figure S3), which also shows that all seven RhoGTPases appear similar. Both the sequence and structural features highlight several similarities. Still, the ratio of GTPases to GEFs in *Entamoeba* (unpublished data from our lab) indicates a complex network of selective GEF activation of GTPases and their connections to functions.

**Figure 1:**
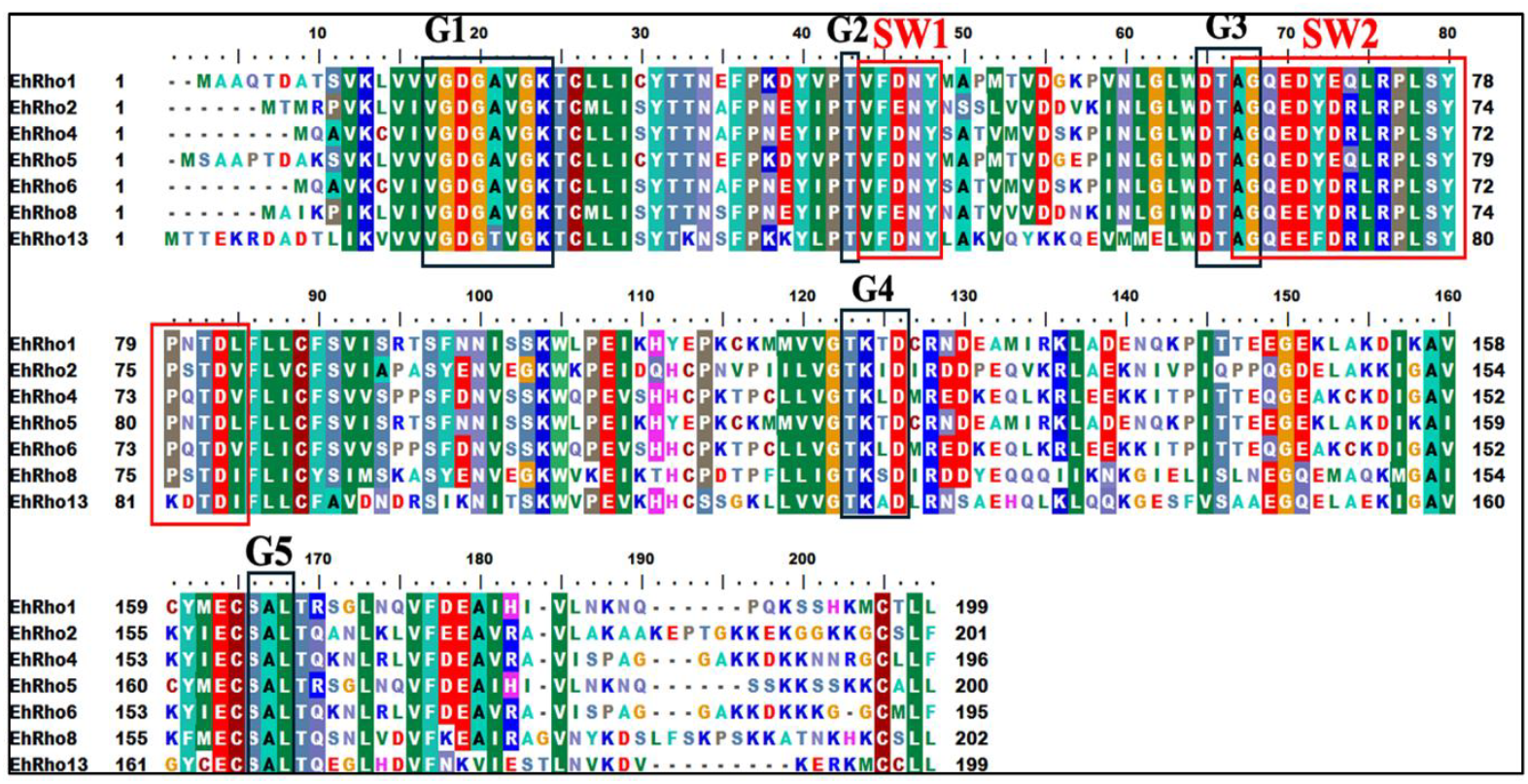
Multiple sequence alignment of the seven Rho GTPase proteins. Conserved regions are highlighted, with boxes indicating the canonical G-box motifs (G1–G5) positions and the Switch I and Switch II regions essential for nucleotide binding and hydrolysis.

### Exchange Activity of GEF (FP10_DH/PH) with Rho GTPases

EhFP10 is a multidomain protein in which the DH-PH domain defines its catalytic function as a GEF. It has no human counterpart, making it highly critical and functionally involved in actomyosin regulation during phagocytosis. The GEF activity of the FP10 protein towards seven selected GTPases (Rho1, Rho2, Rho4, Rho5, Rho6, Rho8, and Rho13) showed catalytic properties at different rates from 3734 to 700 absorbance units/µM (Figure 2). The most pronounced effect was observed with Rho2, which showed a decrease of 3734 units/µM in GTPase activity. Rho6 exhibited the next highest activity with a decrease of absorbance of 2166 units/µM, followed by Rho8 (1300 units/µM), Rho13 (1183 units/µM), Rho4 (1225 units/µM), Rho5 (908 units/µM), and Rho1 (700 units/µM), displaying the lowest activity.

**Figure 2:**
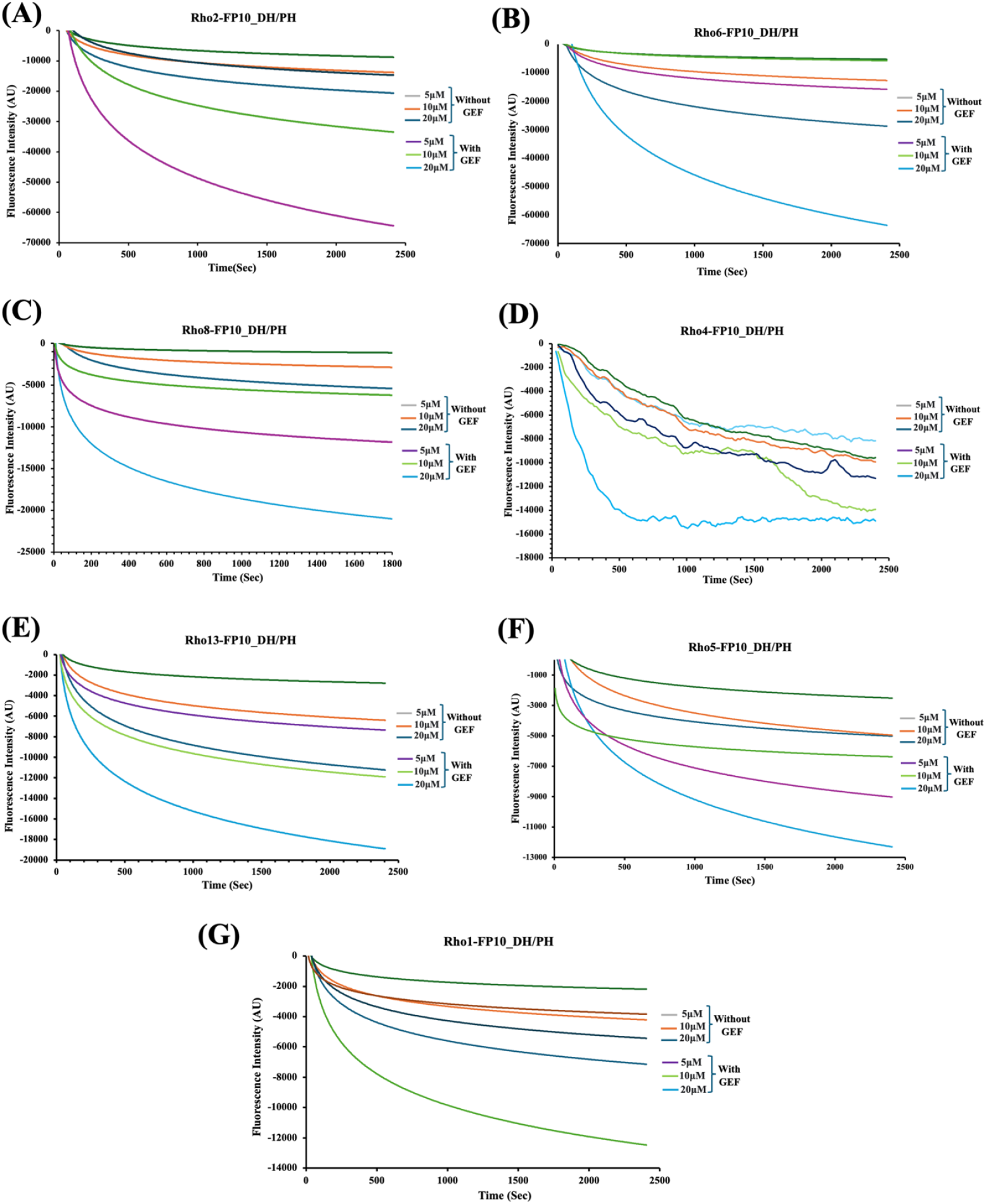
Nucleotide exchange activity of Rho GTPase proteins with FP10_DH/PH domain. The graph shows fluorescence intensity (y-axis) plotted against time (x-axis). Rho GTPases were preloaded with mant-GDP, and a decrease in fluorescence intensity indicates dissociation of mant-GDP, indicating the exchange of GDP for GTP. Among the tested proteins, Rho2 and Rho6 showed the highest exchange rates, followed by moderate activity in Rho8, Rho4, and Rho13. Rho5 and Rho1 displayed the lowest activity.

Based on this observed gradient of GEF activity, the tested GTPases can be categorized into three groups: High Activity: Rho2 & Rho6; Moderate Activity: Rho8, Rho13, and Rho4; and Lower Activity: Rho5 and Rho1. The comparative analysis of the guanine exchange activity of all seven Rho’s, with the control as highest (100%), Rho2, it has been 58% for Rho6, moderately active Rho8 (34.80%), Rho4 (32.80%), Rho13 (31.68%) and lowest Rho5 and Rho1 were 24.30% and 18.74% respectively (Figure 3).

**Figure 3:**
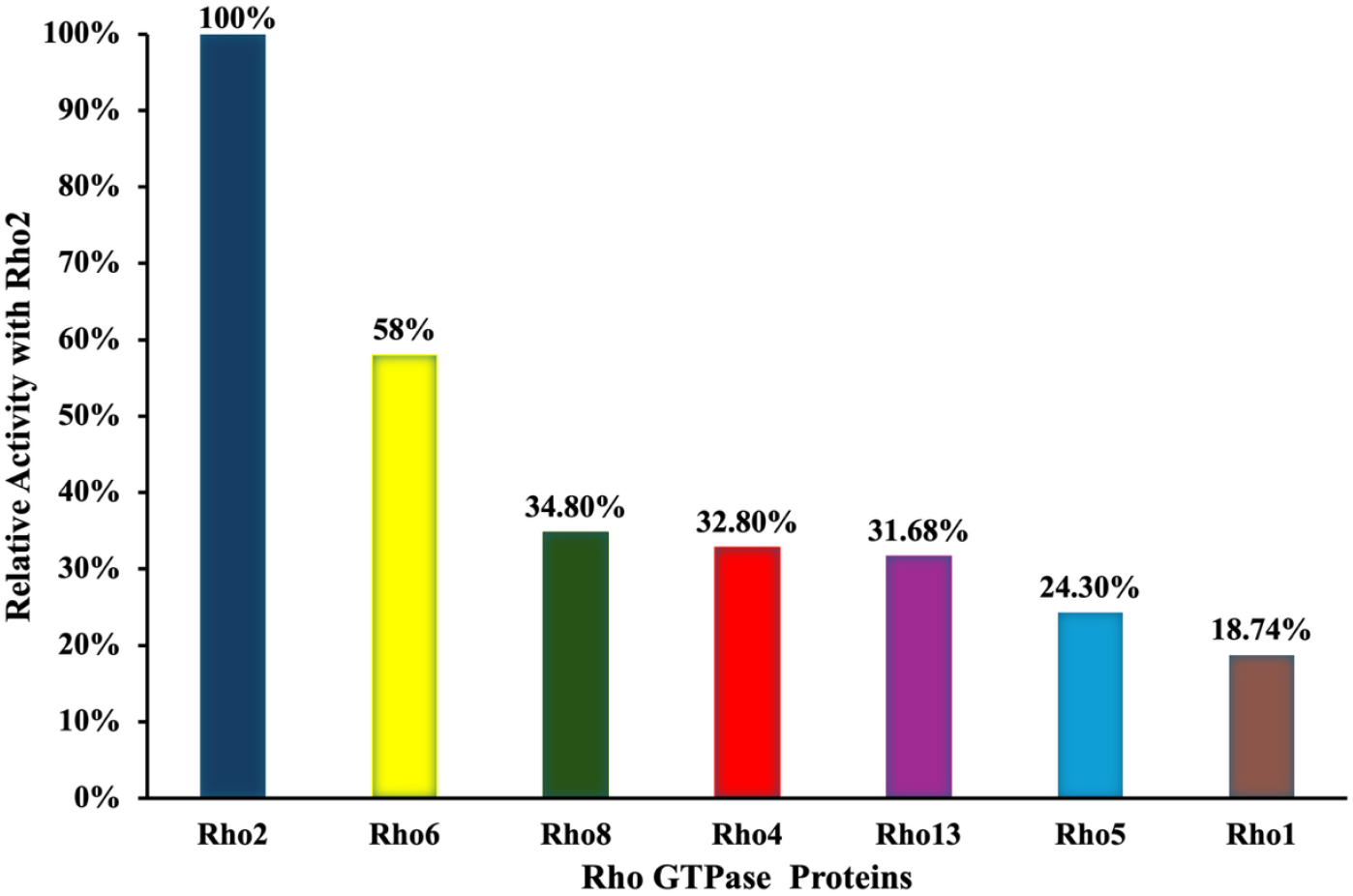
Comparative analysis of GEF-mediated nucleotide exchange activity of six Rho GTPases relative to Rho2. The GTPase proteins were categorized based on their exchange activity with the Rho2 GEF. Blue and yellow indicate high exchange activity. green, red, and purple indicate moderate activity, and cyan and brown represent low activity. This classification highlights the differential responsiveness of Rho GTPases to Rho2-mediated activation.

### Structure-Sequence Analysis of GEF-GTPase complexes

All seven Rhos (Rho1, Rho2, Rho4, Rho5, Rho6, Rho8, and Rho13) share 50-60% sequence identity with each other. The multiple sequence alignment shows that all seven EhRhos share conserved regions, switch regions, G-boxes, and specific residues, but differ in other regions (see Figure 1). Rho2 has a sequence identity of 51.5% with Rho5, as they are in two distinct sets of guanine exchange activity. The individual assessment of conserved residues in the regions responsible for performing the activity is shown in Table 1. Sequence comparison of Rho GTPases reveals a strong correlation between specific residue conservation and GEF-mediated nucleotide exchange activity. Rho2, which exhibits the highest activity, retains critical interacting residues—Glu33, Arg4, and Leu69 —that are consistently absent in lower-activity GTPases such as Rho1, Rho4, and Rho13. Notably, Asn43 in Rho2 is substituted by Ser41 in Rho6 and Rho4, which show slightly reduced but still significant activity, suggesting partial functional compensation. In contrast, Rho1 and Rho5, which exhibit the lowest GEF activity, lack Glu33, Arg4, and Leu69 and show key substitutions including Met (replacing Asn43) and Gln (replacing Arg68).

**Table 1:** Comparison of all seven interacting residues of Rho-GTPase with the GEF (FP10_DH/PH domain) protein.

| Rho2 | Rho6 | Rho8 | Rho4 | Rho13 | Rho5 | Rho1 |
| --- | --- | --- | --- | --- | --- | --- |
| Glu(33) | Glu(31) | Glu(33) |  |  | Asp(38) |  |
| Tyr(34) | Tyr(32) | Tyr(34) | Tyr(32) | Tyr(40) | Tyr(39) | Tyr(38) |
| Thr(37) | Thr(35) | Thr(37) | Thr(35) | Thr(43) |  | Thr(41) |
| Val(38) | Val(36) | Val(38) | Val(36) | Val(44) | Val(38) | Val(42) |
| Arg(4) |  |  |  |  |  |  |
| Asn(43) | Ser(41) | Asn(43) | Ser(41) | Leu(43) | Met(48) | Met(47) |
| Asn(41) | Asn(39) | Asn(41) | Asn(39) | Asn(47) | Asn(46) | Asn(45) |
| Phe(39) | Phe(37) | Phe(39) | Phe(37) | Phe(45) | Phe(44) | Phe(43) |
| Asp(59) | Asp(57) | Asp(59) | Asp(57) | Asp(65) |  | Asp(63) |
| Trp(58) | Trp(56) | Trp(58) | Trp(56) | Trp(64) | Trp(63) | Trp(52) |
| Gln(63) | Gln(61) | Gln(63) | Gln(61) | Gln(69) |  | Gln(67) |
| Leu(69) |  |  |  |  |  |  |
| Arg(68) | Arg(66) | Arg(68) | Arg(66) | Arg(74) | Gln(73) | Gln(72) |
| Tyr(66) | Tyr(64) | Tyr(66) | Tyr(64) |  | Tyr(71) | Tyr(70) |
|  | Ala(59) |  |  |  |  |  |
|  | Asp(38) |  |  |  | Asp(45) | Asp(44) |
|  |  |  |  | Met(60) |  | Asn(58) |
|  |  |  |  | Glu(71) | Asp(70) |  |
|  |  |  |  | Asp(82) |  |  |
|  |  |  |  | Lys(81) |  |  |
|  |  |  |  | Glu(62) |  |  |
|  |  |  |  | Ile(75) |  |  |
|  |  |  |  |  | Glu(72) |  |

Comparative analysis of interacting residues revealed that GTPases with reduced GEF-stimulated activity in Rho4, Rho13, and Rho1 lacked critical switch I contacts, such as Glu33 and Leu69 in Rho2, which mediate electrostatic and hydrophobic interactions with the GEF interface (Figure 4A). In the case of Rho5, Glu is replaced by Asp, which also impedes interface interactions in the switch I region. The missing interactions between both Glu and Leu and GEF (FP10_DH/PH) also corroborate the observation that activity is most severely impaired across all four Rho GTPases. Additionally, substitution of Asn43 of Rho2 with Met in Rho5 and Rho1, which is located immediately C-terminal to Switch I, along with the substitution of Arg 68 of Rho2 with Gln, which lies within Switch II, also contributes to specificity against GEF. The cumulative changes observed in Rho5 and Rho1 are anticipated to significantly hinder GEF engagement at Switch I and disrupt Mg^2+^ stabilization through the GEF acidic finger mechanism at Switch II (Figure 4B). This is a key reason why these proteins exhibit the lowest nucleotide exchange activity. The alterations not only diminish the specificity of the GTPase-GEF(FP10_DH/PH) complex but also compromise its stability. Consequently, the progressive loss or modification of critical amino acid residues of Switch I and Switch II is closely associated with a decline in GTPase activity, underscoring their vital importance in facilitating effective GEF interactions and promoting nucleotide exchange (Table S2).

**Figure 4:**
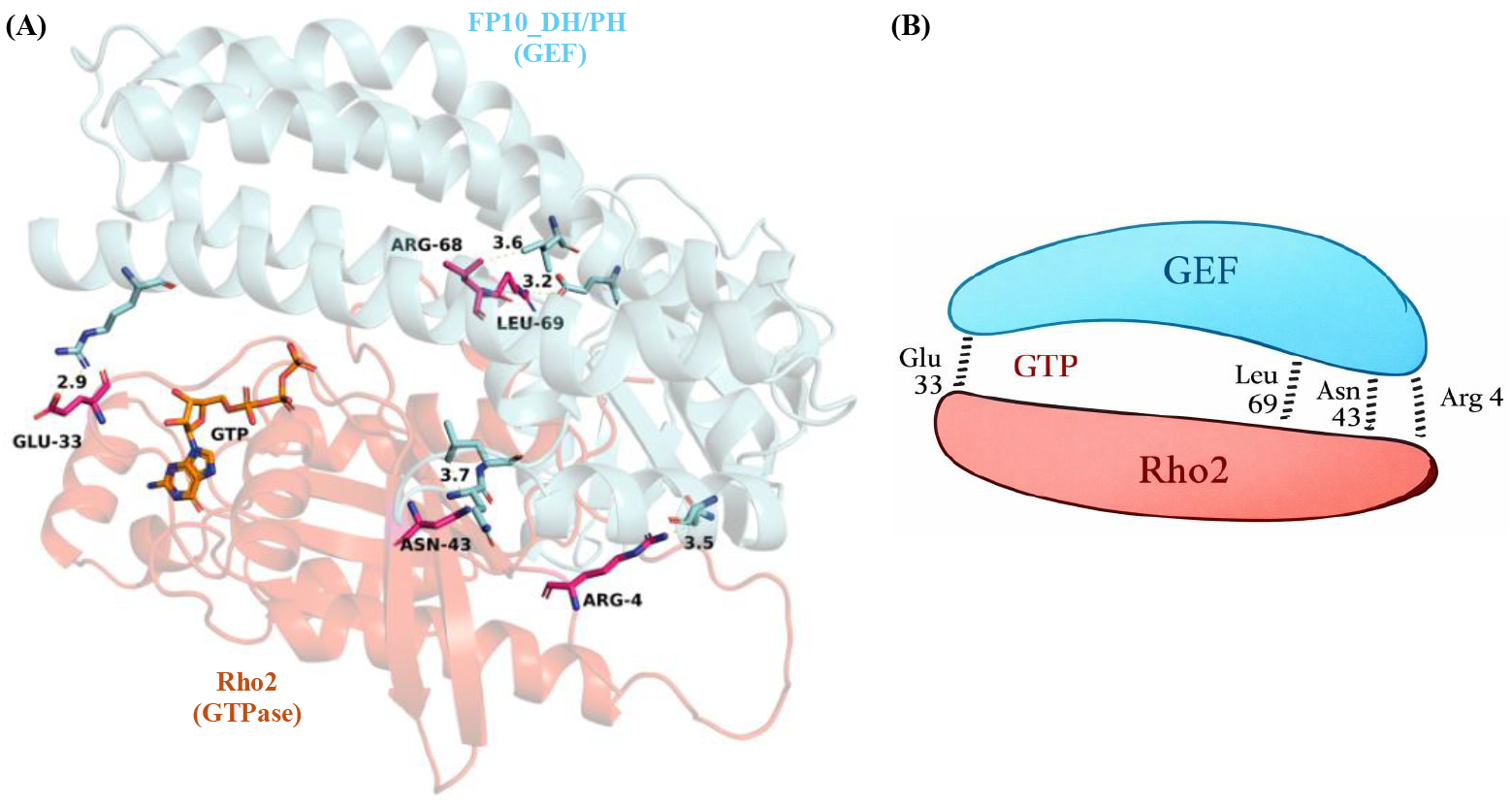
Mechanistic representation of Rho2 and FP10_DH/PH at the residue level involved in GEF exchange in switch 1 and switch 2 regions. (A) Rho2-FP10_DH/PH complex structure model bound to GTP and showing the important residues of Switch I and Switch II involved in interaction during exchange activity. (B) Cartoon diagram showing the open-close mechanism for entering GTP during the transient reaction. Switch I: Missing Glu→Loses initial GEF docking contact. Switch I: Missing Leu→Loses hydrophobic interface stabilization. Post-SW1: Asn→Met→Switch I cannot open (steric block + lost H-bond). Switch II: Arg → Gln → GEF acidic finger cannot insert, Mg^2+^ not destabilized.

In vitro activity and sequence-based analysis were further validated in silico via MD simulations of the GTPase-GEF modeled complex with GTP for all seven Rho GTPases to understand the specificity and stability of the complex (Table 2).

**Table 2:** Comparison of MD simulation parameters of all seven Rho-GTPase-GEF (FP10_DH/PH) protein complexes.

| Complex name | RMSD (nm) | RMSF (nm) | Radius of Gyration (nm) | Surface area (mm/S2/N) | Err. Est | Total Energy (Kj/Mol) | Total Drift (Kj/Mol) |
| --- | --- | --- | --- | --- | --- | --- | --- |
| Rho2-FP10_DH/PH | 0.63±0.14 | 0.24±0.13 | 2.74±0.056 | 276.02±3.88 | 29 | -9.82545e+05 | -76.204 |
| Rho6-FP10_DH/PH | 0.43±0.11 | 0.26±0.10 | 2.70±0.0541 | 261.48±4.18 | 73 | -1.19052e+06 | -505.844 |
| Rho8-FP10_DH/PH | 0.37±0.11 | 0.26±0.12 | 2.64±0.047 | 267.45±5.41 | 23 | -1.02767e+06 | -150.784 |
| Rho4-FP10_DH/PH | 0.49±0.12 | 0.28±0.13 | 2.70±0.069 | 273.14±4.80 | 50 | -1.00481e+06 | -362.098 |
| Rho13-FP10_DH/PH | 0.47±0.08 | 0.17±0.09 | 2.53±0.035 | 265.74±5.55 | 55 | -1.19837e+06 | -372.098 |
| Rho5-FP10_DH/PH | 0.56±0.17 | 0.53±0.15 | 4.72±0.0643 | 269.42±4.85 | 110 | -1.19021e+06 | -499.247 |
| Rho1-FP10_DH/PH | 1.71±0.68 | 2.33±1.16 | 4.87±0.10 | 280.19±4.41 | 36 | -1.23616e+06 | -254.348 |

The root mean square deviation (RMSD) was calculated to evaluate the stability and flexibility of the Rho–FP10_DH/PH complexes (Figure 5A). The Rho2–FP10_DH/PH complex demonstrated an average RMSD of approximately 0.68 nm, suggesting a flexible and dynamic structure conducive to nucleotide exchange. In contrast, Rho6, Rho8, Rho4, and Rho13 exhibited lower RMSD values compared to Rho2, indicating more compact and stable complexes with moderate GEF activity. RMSD values exceeding 0.8 nm were deemed indicative of structural instability. The Rho1 complex showed an average RMSD of 1.71 nm, with some regions exceeding 2.05 nm, consistent with its low activity. Meanwhile, the Rho5 complex recorded an RMSD of approximately 0.56 nm but demonstrated diminished activity, likely due to variations in residue interactions.

**Figure 5:**
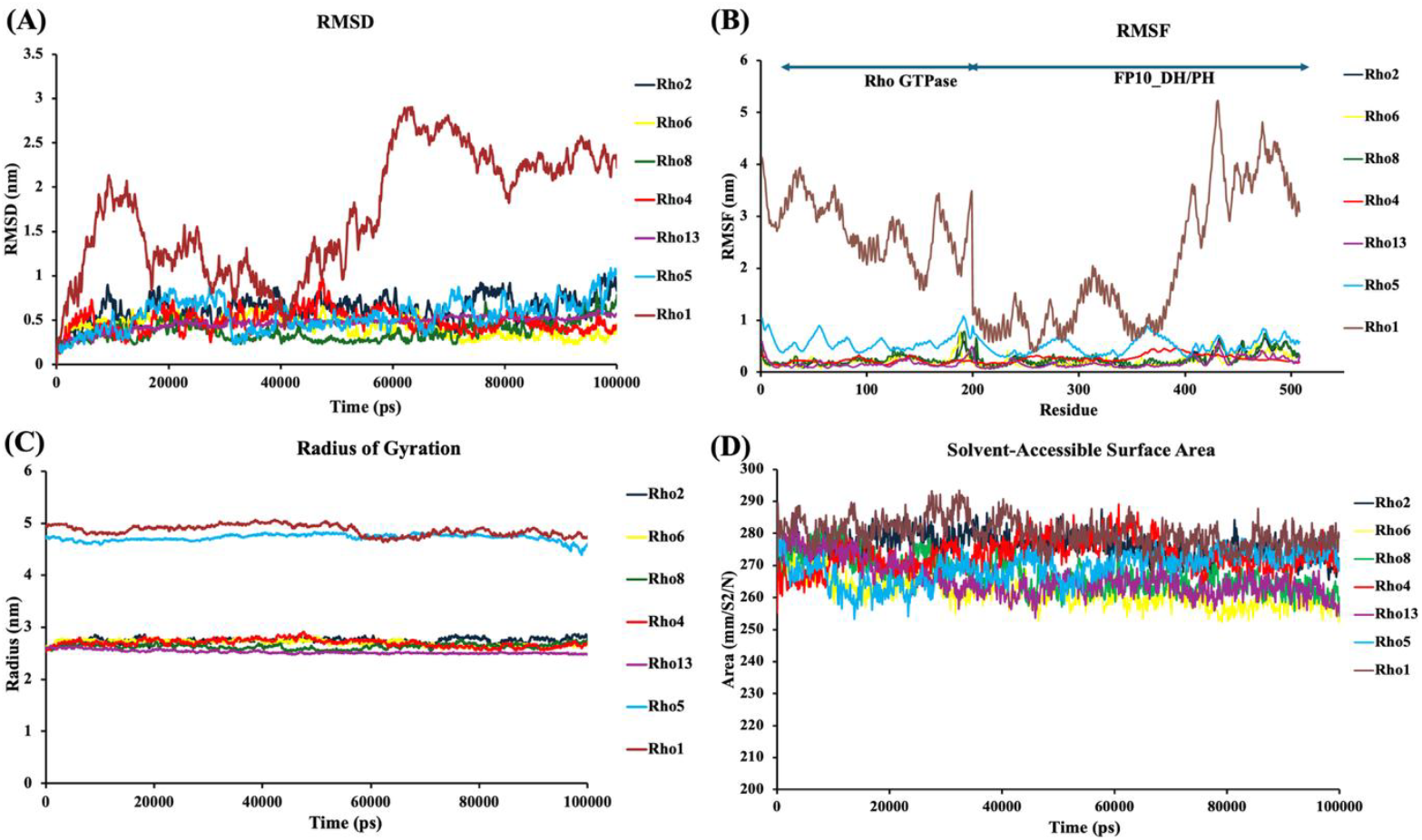
MD simulation of all seven Rho GTPase-GEF (FP10_DH/PH) Complexes. A) Backbone RMSD profiles of all seven complexes over the 100 ns simulation period, illustrating the overall structural stability and equilibration behavior of each system. Most complexes exhibit stable trajectories with minor fluctuations, whereas the Rho1-FP10_DH/PH complex shows comparatively higher deviation, indicating reduced conformational stability. (B) RMSF per residue, highlighting local flexibility across the protein sequence. Elevated fluctuations are primarily observed in loop regions, while the DH and PH core domains remain relatively stable, consistent with their functional rigidity. (C) Rg plots demonstrating the compactness of each complex throughout the simulation. The relatively constant Rg values indicate preservation of structural integrity, with no significant unfolding events. Rho1 and Rho5 complexes with GEF show higher values and are distant during the simulation period. (D) SASA analysis showing the extent of solvent exposure during the trajectory. The minor variations in SASA suggest stable folding and consistent surface accessibility across all complexes.

**Figure 6:**
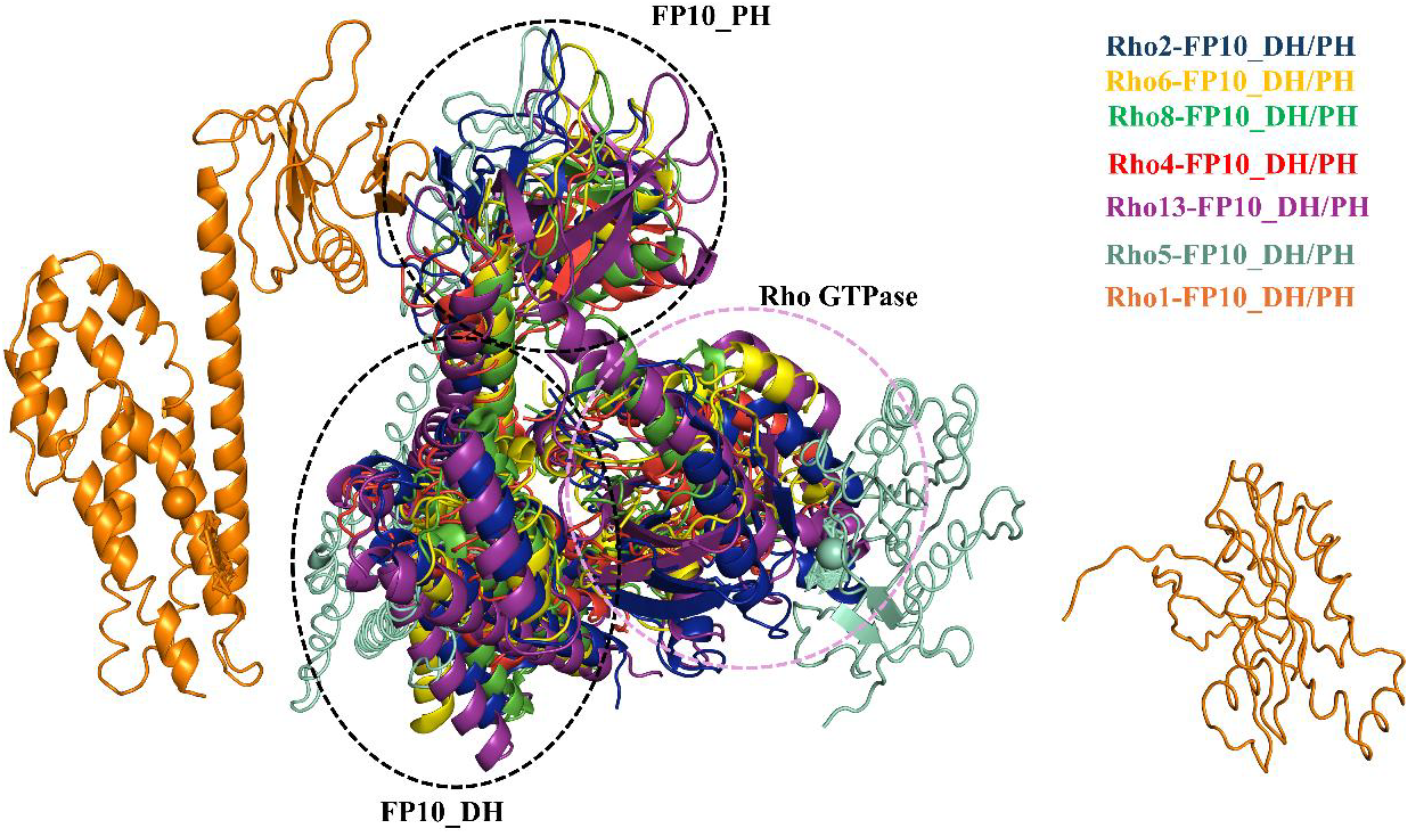
Structural Comparison of all seven Rho GTPase-GEF (FP10_DH/PH) Complexes. Superimposed structures of the average PDBs from MD simulations of all seven Rho GTPase–GEF (FP10_DH/PH) complexes are shown, highlighting their overall structural alignment and conformational similarities. Each complex is represented in a distinct colour as indicated in the legend. The Rho GTPase is centrally positioned, while the FP10 DH and PH domains are indicated and circled for Rho2, Rho6, Rho8, Rho4, and Rho13; Rho5 and Rho1 are out of place in complex formation.

Root mean square fluctuation (RMSF) was analyzed for the conserved G motifs (G1–G5) of Rho GTPases (Figure 5B). In the Rho2–FP10_DH/PH complex, residues 15–20 (G1), 45–50 (G2/switch I), 70–80 (G3/switch II), 120–125 (G4), and 165–170 (G5) displayed low fluctuations (<0.25 nm), indicating a rigid nucleotide-binding cage stabilized by FP10_DH/PH. Rho1 exhibited pronounced fluctuations at G2/G3 and moderate fluctuations at G4–G5, with values up to 2.33 nm, reflecting instability in the catalytic domain. Rho5 showed fluctuations of ~0.53 nm, accompanied by higher RMSD and Rg values, consistent with weak stability. Rho6, Rho4, Rho8, and Rho13 displayed fluctuation patterns similar to Rho2, correlating with moderate activity.

The radius of gyration was measured to evaluate protein compactness (Figure 5C). For complexes of ~510 residues, Rg values between 2.8 and 3.4 nm indicated stable, globular structures. Rho2, Rho6, Rho8, Rho4, and Rho13 maintained Rg values below 3 nm throughout the 100 ns simulation, confirming stable folding. In contrast, Rho1 and Rho5 exhibited Rg values around 5 nm, suggesting unstable or unfolded complexes.

SASA values for all Rho–FP10_DH/PH complexes were approximately 275 nm^2^, indicating comparable solvent exposure (Figure 5D). The total energy of the complexes averaged ~1.0 × 10^−6^. Energy drift analysis revealed that the Rho2–FP10_DH/PH complex had the lowest drift energy (–76.0 kJ/mol), highlighting its superior stability compared to the other complexes.

The structural analysis of the Rho-FP10_DH/PH complexes, compared with their simulated average structures, reveals that the Rho2-FP10_DH/PH complex exhibits moderate conformational similarity with other Rho proteins, such as Rho6, Rho8, Rho4, and Rho13. In contrast, the average structures of the Rho1-FP10_DH/PH and Rho5-FP10_DH/PH complexes do not align well with the Rho2-FP10_DH/PH complex. This discrepancy suggests that Rho1-FP10_DH/PH and Rho5-FP10_DH/PH exhibit weaker sustained interactions or lower affinity for the FP10_DH/PH component, as indicated by their structural separation from the Rho2 complex.

To investigate the structural basis of differential GEF activation among Rho GTPases, LigPlot+ interaction analysis was performed on modeled complexes between each Rho isoform and the DH domain of the FP10 protein. The two-dimensional plots visualize hydrogen bonding and hydrophobic contacts, revealing distinct interaction patterns correlated with GTPase activity (Figure S4). Rho2– FP10_DH/PH is the most active complex, exhibiting an extensive interaction network involving Glu33, Arg4, Leu69, Asn43, and Arg68, supported by both central and peripheral hydrogen bonds and hydrophobic contacts. In contrast, GTPases with moderate activity, such as Rho6, Rho8, and Rho4, showed partial deviations, including Asn43→Ser41 and Arg68→Gln72, or a complete absence of Glu33 and Arg4, resulting in weakened interactions at key sites. In Rho13 and Rho5, the LigPlot+ diagrams revealed additional substitutions, such as Met48 and Glu73, and further loss of native hydrogen bonds at the Asn133 and Arg150 interaction sites, corresponding to a further drop in activity. The Rho1–FP10_DH /PH complex, with the lowest GEF activity, showed the most disrupted interface: contact losses at nearly all key residues (e.g., Glu33, Arg4, Leu69) and widespread substitutions (Met47, Gln72), leading to fragmentation of the interaction network. These differences are visualized in the circled and boxed regions across the plots, with progressively fewer interaction lines and reduced bonding density from Rho2 to Rho1. The LigPlot+ data demonstrate a direct correlation between contact conservation and functional activation. Isoforms that retain Rho2-like interaction motifs form tighter, more stable GEF complexes, while deviations at critical positions weaken binding and reduce GEF activity. These 2D representations reinforce the hypothesis that specific residue-level interactions are essential for productive GTPase-GEF engagement, validating the functional significance of the identified contact residues.

To investigate inter-residue dynamic behavior during Rho–FP10_DH/PH domain complex formation, dynamic cross-correlation analysis (DCCM) was performed across all isoforms. (Figure S5) These matrices show the extent to which the movement of residues is correlated (red, positive), anti-correlated (blue), or uncorrelated (white). It provides insight into concerted motions and functional relationships between different protein regions. These findings collectively suggest that optimal interaction with GEF (FP10_DH domain) requires a balance of rigidity and localized flexibility. Rho2–FP10_DH/PH, Rho6– FP10_DH/PH, and Rho8– FP10_DH/PH complexes show dominant red regions, reflecting strong positive correlations between residues. In contrast, the Rho4–FP10_DH/PH, Rho5– FP10_DH/PH, Rho13–FP10_DH/PH, and Rho1–FP10_DH/PH complexes show reduced red colouration, indicating weaker or less correlated residue motions. The overall red intensity follows a gradient from high in the Rho2–FP10_DH/PH complex to low in the Rho1–FP10_DH/PH complex. The low deformability of key interacting residues in the Rho2–GEF complex supports strong, stable binding. At the same time, the more flexible or poorly structured interfaces in low-activity GTPases reduce interaction fidelity and GEF-mediated activation.

The results show a strong relationship between coordinated residue motion and GTPase activity. The Rho2–FP10_DH domain complex exhibits the highest GEF activity and shows broad and intense red regions extending across diagonals and off-diagonal zones. This indicates highly coordinated motion between distant residues across the Rho and GEF domains, reflecting a tightly coupled and functionally optimized interface. Similarly, Rho6 and Rho8, with moderately high activity, also display substantial correlated motion but slightly less uniform red distribution, suggesting minor reductions in interface coupling efficiency. In contrast, lower-activity complexes such as Rho4, Rho5, and Rho13 exhibit patchier and more fragmented red zones, with more blue (anti-correlated) and white (non-correlated) regions, reflecting weaker intramolecular communication. The Rho1–FP10_DH/PH domain complex, consistent with its lowest GEF activity, shows the most scattered and disrupted correlation pattern, with large uncoordinated areas and weak or absent long-range coupling. This result indicates that the absence of residue-residue communication dynamics in Rho1 likely contributes to inefficient GEF interaction and impaired catalytic function.

### Specific interaction of GEF (FP10_DH/PH Domain) with Rho GTPases

The activity of the guanine nucleotide exchange factor with various GTPases suggests that this GEF (FP10_DH/PH) can activate multiple GTPases in a gradient manner. We focused on three selected GTPases representing a gradient of activity: Rho2 (high), Rho4 (moderate), and Rho1 (low) to analyze interactions between complexes using Bio-Layer Interferometry (BLI). The interaction analysis revealed that the FP10_DH/PH complex showed the highest binding affinity for Rho2, with a dissociation constant (KD) of 0.58 µM. In contrast, Rho4 and Rho1 exhibited KD values of 1.3 µM and 1.9 µM, respectively (Figure 7). Notably, the high exchange activity is correlated with increased affinity, as demonstrated by complementary protein-protein interaction data. Moreover, the lower binding affinities observed for Rho4 and Rho1 suggest distinct kinetic profiles, reflecting their diminished exchange capabilities. These findings highlight the importance of transient interactions for nucleotide exchange mediated by GEFs.

**Figure 7:**
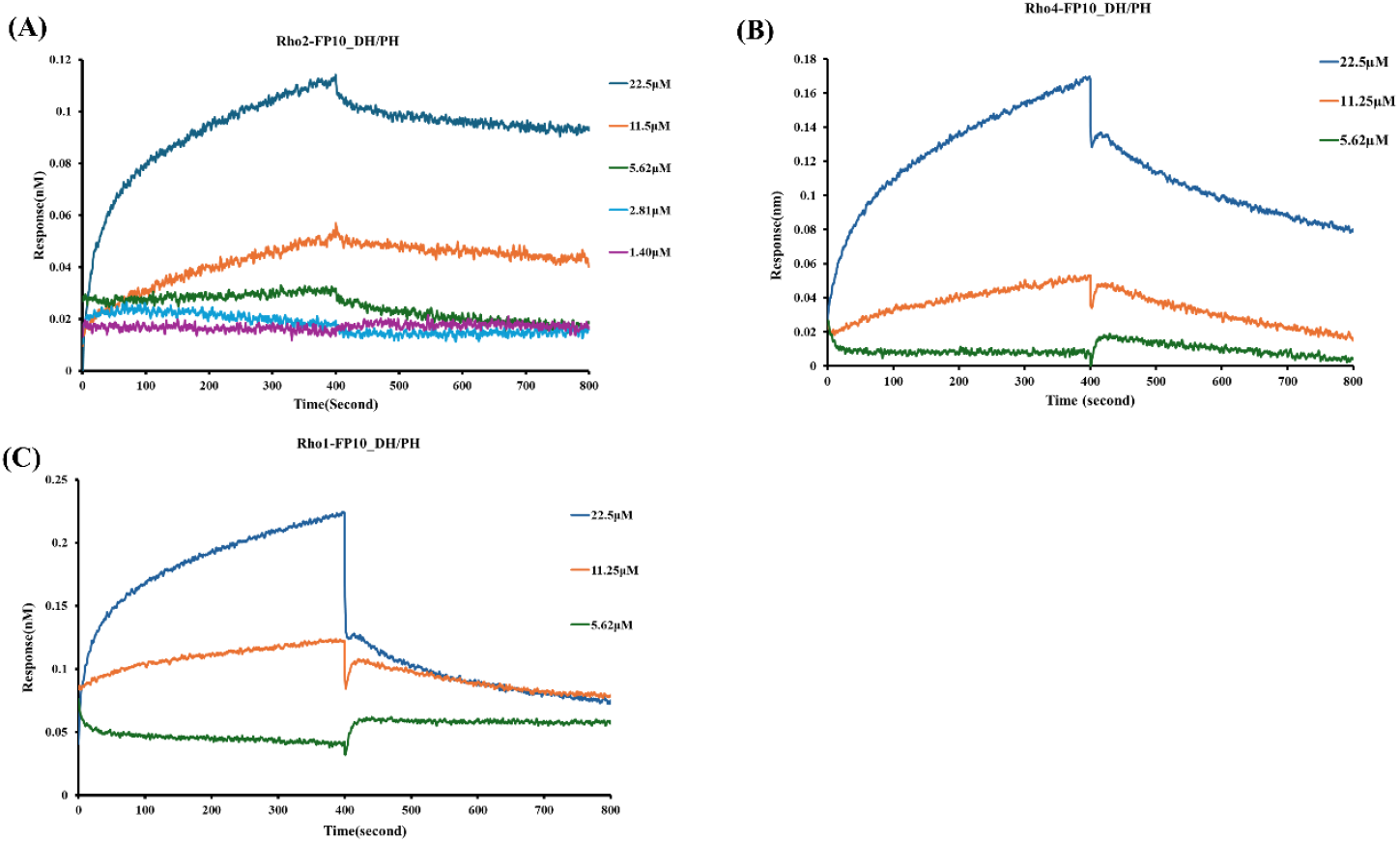
Binding analysis of Rho GTPase proteins with FP10_DH/PH. Biolayer Interferometry (BLI) analysis shows interactions between Rho2, Rho4, and Rho1 and the FP10_DH/PH (GEF) protein. The binding response (y-axis) is plotted against time (x-axis), with the first 400 seconds representing the association phase and the following 400 seconds the dissociation phase. Binding affinities (K_D_) were calculated from these curves: Rho2–GEF (FP10_DH-PH domain) = 0.58 µM, Rho4–GEF = 1.3 µM, and Rho1–GEF = 1.9 µM.

The activity and protein-protein interaction indicate that the affinity of the protein is directly related to the specificity of the GTPase and GEF in that particular event. However, in addition to specific interactions, other Rho GTPases also compensate for the exchange process, albeit with less specificity or in the absence of the particular molecule.

## DISCUSSION

All seven Rho GTPase proteins (Rho1, Rho2, Rho4, Rho5, Rho6, Rho8, and Rho13) and FP10 have already been reported in the literature to be involved in signaling pathways for endocytic processes such as phagocytosis, pinocytosis, and trogocytosis ^27,31–34^. These GTPases function as molecular switches, undergoing conformational transitions between inactive GDP-bound and active GTP-bound states, driven by Guanine nucleotide Exchange Factors (GEFs), most notably those of the Dbl family ^28,35–37,40– 43^. EhFP10 is one of the GEFs with an unusual domain architecture, only seen in amoebic species, and is involved in Actin and myosin binding and phagocytosis.

In this study, we systematically investigated the interaction between the GEF domain of FP10 and a panel of seven Rho family GTPases from *E. histolytica* to identify the specificity and functional regulation. If one GEF is around several Rho GTPases, can one GEF interact with all Rho GTPases, or does it have any specificity? What regions of the GEF determine the specificity?

Using a combination of GEF activity assays, protein-binding studies, and molecular dynamics simulations, the data consistently identified Rho2 as the most specific and high-affinity interactor of FP10. This suggests the existence of a selective GEF–GTPase signalling axis in *E. histolytica*, akin to mammalian RhoGEF-RhoGTPase modules ^44–47^. The structural determinants underpinning this specificity were further revealed through sequence and interaction profiling. Rho2 retained critical contact residues such as Glu33, Arg4, Leu69, and Asn43, which were progressively lost or substituted in other isoforms, particularly those with lower GEF responsiveness (e.g., Rho1, Rho5, and Rho13). Conformational shifts in response to nucleotide exchange are dominated by two conserved switch regions that contribute to nucleotide state-selective engagement of multiple Rho-interacting proteins. Switch 1 contains a highly conserved phenylalanine that forms aromatic interactions with the nucleotide guanine ring. The importance of CAAX box-mediated posttranslational modifications, including isoprenylation, for subcellular localization and function of these GTPases has also been noted ^44,48,49^.

The absence of Glu and Leu from Switch I likely disrupts the formation of the initial recognition complex between the GTPase and GEF, consistent with structural studies indicating that these residues contribute to the buried interface. The Asn-to-Met substitution immediately after Switch I introduces steric incompatibility and eliminates a hydrogen bond that is crucial for stabilizing the open Switch I conformation required for GDP egress. Additionally, the Arg-to-Gln substitution in Switch II removes the positive charge needed to properly orient Switch II during the insertion of the GEF acidic finger, thereby impairing Mg^2+^ dissociation, which is essential for nucleotide exchange. The convergence of these defects across both switch regions offers a structural rationale for the nearly complete loss of GEF-stimulated GTPase activity observed in these variants.

While such specificity has been well characterized in higher eukaryotes ^47,49–52^, its molecular basis remains unexplored, particularly in protozoan parasites. The present study’s findings suggest that FP10 may preferentially activate Rho2 through a selective recognition mechanism. This interaction likely plays a pivotal role in modulating actin cytoskeleton remodeling and endocytic processes, essential for *E. histolytica’s* motility, phagocytosis, and overall pathogenicity. Molecular dynamics simulations provided additional insights into the stability and functional compatibility of these complexes. The Rho2–FP10_DH/PH complex displayed minimal deformability and extensive correlated residue motions—characteristics of a well-coupled protein-protein interface necessary for efficient signal transmission ^52,53^. Conversely, less active complexes (e.g., Rho1–FP10_DH/PH) showed reduced coordination and interaction stability, reflecting impaired catalytic efficiency. These *in vitro* data provide compelling support for the specificity and functional relevance of the FP10–Rho2 interaction. LigPlot+ interaction diagrams showed that high-activity complexes exhibited denser hydrogen bonding and hydrophobic interactions, whereas lower-activity complexes exhibited fragmented and weakened interfaces.

The convergence of evidence across multiple methodologies underscores the significance of this interaction, demonstrating high-affinity binding between FP10 and Rho2, corroborated by increased nucleotide exchange rates in GEF assays. These findings demonstrate that the interaction between FP10 and Rho2 is highly selective, structurally reinforced, and functionally robust. This isoform-specific GEF-GTPase pairing potentially orchestrates key cytoskeletal rearrangements that underpin the parasite’s invasive processes. Although this selectivity mirrors mechanisms in higher eukaryotes, ^47,49,52,54,55^ it is newly described in protozoan pathogens and offers a paradigm for understanding signal transduction specificity in *E. histolytica*. The modular architecture of the FP10 GEF—comprising a DH domain for catalytic activity and a PH domain likely involved in membrane recruitment and spatial guidance ^45,56^—supports a model wherein Rho2 is activated at specific membrane locales during cytoskeletal remodeling.

The molecular dynamics simulations and structural comparisons demonstrate that the stability and activity of Rho–FP10_DH/PH complexes are strongly influenced by the interacting residues at the binding interface. The Rho2–FP10_DH/PH complex showed moderate conformational similarity with Rho6, Rho8, Rho4, and Rho13, which share conserved residues that maintain stable contacts with FP10_DH/PH. This conservation supports compact structures, balanced flexibility, and moderate GEF activity, consistent with their RMSD, RMSF, and Rg values. In contrast, Rho1 and Rho5 showed distinct differences in their interacting residues, resulting in poor alignment with Rho2 and reduced stability. Substitutions at critical positions disrupted sustained interactions with FP10_DH/PH, leading to higher RMSD and Rg values, greater fluctuations in the G motifs, and overall structural instability. These residue-level differences explain the weak affinity and minimal GEF activity observed for Rho1 and Rho5. Together, the findings highlight that conserved interacting residues are essential for maintaining productive conformations and efficient nucleotide exchange, while divergence at these positions compromises complex stability and function.

### Conclusion

Conclusively, this study identifies a novel and selective regulatory axis between the EhFP10 (GEF) and EhRho2 in *E. histolytica*, suggesting it may serve as a central modulator of actin dynamics and pathogenesis. This interaction, now structurally and biochemically defined, represents a promising therapeutic target. The specificity of selecting for RhoGTPase against a GEF depends on its physiological role, as seen among the seven RhoGTPases studied, given that their switch regions are conserved and crucial for exchange activity. Also, the pattern of protein-protein interactions and complex model simulations indicates that, due to transient interactions, optimal interactions are more important than strong or weak binding for this signaling axis, which is associated with several pathogenic functions.

### Future Perspective

Future in vivo studies involving gene manipulation (e.g., knockdown or overexpression of Rho2 with FP10) will be crucial to elucidate the physiological relevance of this axis. Additionally, identifying downstream effectors of Rho2 would further delineate the signaling network regulating cytoskeletal architecture in this parasite.

## MATERIAL & METHODS

### Expression and purification of Rho GTPase proteins

The seven proteins of interest (Rho1, Rho2, Rho4, Rho5, Rho6, Rho8, and Rho13) were purified using Ni-NTA (nickel-nitrilotriacetic acid) affinity chromatography^57^. *E. coli* C41(DE3) competent cells were transformed with the respective clones and cultured until an optical density (OD) of 0.6-0.7 was reached at 600 nm. Protein expression was induced using isopropyl β-D-1-thiogalactopyranoside (IPTG) at a final concentration of 0.25 mM, followed by incubation at 16°C for 12-15 hours. After induction, the cultures were harvested via centrifugation at 8000 rpm for 5 minutes. The resultant cell pellet was resuspended in lysis buffer. A freeze-thaw method was performed for 4-5 cycles to achieve effective cell lysis, resulting in a thicker, more viscous lysate. The viscosity was further reduced by applying sonication (15 sec ON – 20 sec OFF cycle for 15 cycles, amplitude – 40%) to the lysed cell suspension, followed by centrifugation at 18000 rpm for 30 minutes to clear the lysate. The supernatant was then collected and applied to a pre-equilibrated Ni-NTA column using a suspension/equilibration buffer (50 mM Tris-Cl pH 8.0, 150 mM NaCl, 3 mM DTT, 20 mM Arginine, 5% Glycerol) to facilitate binding. The His-tagged proteins bound to the Ni-NTA beads, while unbound proteins were removed by washing the column with 10-15 mL of wash buffer (50 mM Tris-Cl pH 8.0, 150 mM NaCl, 3 mM DTT, 20 mM Arginine, 5% Glycerol, 10 mM Imidazole). The target proteins were eluted using an elution buffer containing a gradient of high concentration of imidazole (100-200 mM), which competes for binding sites on the Ni-NTA, facilitating the release of the His-tagged proteins. Eluted fractions were collected and analyzed, along with the flow-through and wash fractions, by 12% SDS-PAGE to verify purification efficiency and protein yield.

### Mant-GDP fluorescence GEF assay

GEF activity was assessed using a fluorescence-based assay involving mant-GDP (N-Methyl anthraniloyl guanosine diphosphate) ^58,59^. The mant tag, introduced at the ribose moiety’s 2′ or 3′ position in GDP, facilitates real-time monitoring of nucleotide exchange by increasing fluorescence upon protein binding, making it a valuable probe for GTPase interactions ^60^. In our assay, the Rho GTPases were incubated with mant-GDP in a loading buffer comprising 20 mM Tris (pH 7.5), 50 mM NaCl, 0.5 mM MgCl_2_, 10 mM EDTA, 2 mM DTT, and a 10-fold excess of mant-GDP. The incubation proceeded for 4-5 hours at 20°C, protected from light to prevent photodegradation of the fluorescent tag. The reaction was halted by adding 10 mM MgCl_2_ to the mixture. To eliminate unbound mant-GDP, the reaction buffer was exchanged using a Centricon/Amicon device, switching to a buffer containing 40 mM Tris (pH 7.5), 50 mM NaCl, 10 mM MgCl_2_, and 2 mM DTT. Following this purification step, we quantified the protein concentration. Subsequently, the reaction was prepared at 100 µL with varying concentrations of Rho GTPase proteins (20 µM, 10 µM, and 5 µM) in the presence or absence of the GEF (FP10_DH/PH domain) at a fixed concentration of 100 µM. GTP was then introduced to the reaction mixtures. At this point, fluorescence was measured using a spectrofluorometer set to 360 nm excitation and 440 nm emission, with orbital shaking at 282 cycles per minute. A significant increase in fluorescence was observed upon adding GTP in the presence of the FP10_DH-PH domain, indicating that the FP10 accelerates nucleotide exchange, effectively replacing mant-GDP with GTP. This observation was corroborated across all seven Rho GTPases (Rho1, Rho2, Rho4, Rho5, Rho6, Rho8, and Rho13) at identical concentrations, thereby facilitating the comparative analysis of GEF specificity with FP10.

### Bio-layer interferometry

Sartorius’ two-channel system, Octate R2 and Amine Reactive 2nd Generation (AR2G) Biosensor, was used for BLI. The Biosensor (AR2G) was activated by EDC (1-ethyl 3-(3-dimethylaminopropyl) carbodiimide hydrochloride)/ sulfo-NHS (N-hydroxysulfosuccinimide) at optimum concentration. After activation, the FP10_GEF/DH-PH protein was immobilized with a covalent amide bond. The remaining unbound binding site was quenched with 1 M ethanolamine. The other Rho-GTPases, at concentrations ranging from nanomolar to micromolar, were placed in separate wells of the titration plate (black 96-well plate) for the binding assay. Association and Dissociation were set to 300 and 400 seconds, respectively, and the graph shows the binding response over time for each concentration of the other protein. Protein. The K_D_ value was calculated from the Association constant and dissociation constant.

### Structure prediction & protein complex modeling

The protein structure of six Rho-GTPases is not available except for Rho1. The structures of six RhoGTPases (Rho2, Rho4, Rho5, Rho6, Rho8, Rho13) were predicted using the AlphaFold 3.0 web server to understand atomic-level interactions at the structural level^61^. Further, for complex studies, all seven RhoGTPase–GEF (FP10_DH/PH) complexes were modeled individually using AlphaFold3.0 (ColabFold/AlphaFold-Multimer interface). AlphaFold-Multimer, an extension of AlphaFold, predicts the structures of protein complexes by modeling inter-chain contacts and co-evolutionary signals. The target sequences of *E. histolytica* Rho proteins (Rho1, Rho2, Rho4, Rho5, Rho6, Rho8, and Rho13) and the catalytic domains of their cognate GEFs (Dbl homology (DH) and Pleckstrin homology (PH) domains) were retrieved from the UniProt database^62^. The sequences were formatted in FASTA, uploaded to AlphaFold-Multimer, and modeled using default pipeline parameters (MSA construction, template search, structure prediction, and relaxation with the AMBER force field). For each complex, five models were generated, and the best-ranked structure was selected based on predicted Local Distance Difference Test (pLDDT) and predicted aligned error (pAE) scores. Models were visually inspected for interface geometry and domain orientation in PyMOL^63^. The complex structure information was verified using the PDBsum online platform to ensure the reliability of structure models^64^.

### In silico study methods

All selected RhoGTPase sequences were aligned using Clustal Omega ^65^, and images were generated using BioEdit ^66^. All seven Rho GTPase_ GEF (FP10_DH/PH domain) complex 2D Ligplot generated from Ligplot+ software^67^. The iMODS server was used for covariance analysis in the RhoGTPase-GEF complex models^68^. A model covariance map (often called a residue cross-correlation matrix or covariance matrix) from molecular dynamics (MD) simulation results is a visual representation of how the atomic or residue motions are correlated over time. It provides insight into concerted motions and functional relationships between different regions of a protein. The graph shows three colors: Red, Blue, and White. Each colour represents correlation. The red colour indicates that the strongly correlated motions (e.g., domain movements) may indicate allosteric communication. The blue color indicates Anti-correlated regions, which may hint at hinge motions or conformational switches, and the white color indicates non-correlated residue movements in the complex.

### Molecular Dynamics Simulation of the Protein Complexes

Molecular dynamics (MD) simulations were performed on the optimal complex structure using GROMACS ^69^ to assess the binding stability of the protein complexes, with the AMBER99SB-1LDN force field applied to each complex ^70,71^. The MD simulation process concluded after three phases: (i) neutralization of the system, (ii) energy minimization of the system, and (iii) system equilibration. Each protein complex was immersed in a cubic box of 1.0 nanometer (nm) filled with the spc216 water model, and sodium and chloride ions were introduced to neutralize the system during energy minimization. The steepest descent and conjugate gradient algorithms were employed for the energy minimization phase, maintaining a maximum force below 1000 kJ/mol over 50,000 steps. Equilibration of the system was carried out for 50 picoseconds, using the Particle Mesh Ewald (PME) method for electrostatics calculations, which helps produce reliable energy estimates^72,73^. Following system equilibration at a steady temperature of 300 Kelvin, a 100-nanosecond MD simulation was executed every 100 ps. The trajectory files for Root Mean Squared Deviation (RMSD), radius of gyration (Rg), hydrogen bonds, and distances were analyzed to evaluate the binding affinity among the protein complexes. To identify conformational alterations, structural stability, and protein-protein interactions (PPI), both pre- and post-MD structures were superimposed. Various tools were employed to analyze the results of the MD simulation, including Visual Molecular Dynamics (VMD)^74^ for visualizing and animating MD simulation trajectories with 3D graphics, XMGRACE^75^ for plotting data related to RMSD, Rg, hydrogen bonds, and distances, and PyMOL for visualizing the 3D structures extracted from the simulation post-MD. To compute the Solvent Accessible Surface Area (SASA) of the protein complex over the simulation duration, tools such as gmx sasa in GROMACS were used. Changes in SASA were analyzed to understand how solvent-accessible surface area fluctuates during complex formation and dissociation. A reduction in SASA usually signifies stable complex formation, whereas an increase may indicate unfolding or conformational changes that reveal previously buried regions.

## AUTHOR’S CONTRIBUTION

SG, AKG, and PU conceptualized the study. AKG and PU have performed experiments, execution, and data analysis. AKG and PU drafted the initial version of the manuscript, while all the authors critically reviewed and revised the final draft. All authors read and approved the final version of the manuscript. SG generated funds and supervised all the studies.

## ACKNOWLEDGEMENT

The authors gratefully acknowledge DST-FIST-II (grant no. SR/FST/LSII-046/2016(C)), DST-PURSE, and DBT-BUILDER (grant no. BT/INF/22/SP45382/2022) for the Central Instrumentation Facility at JNU and for extending institutional funding. We thank Dr. Anil Raj Narooka and Prof. Sunando Datta from IISER Bhopal for providing all Rho GTPase clones. We also thank Dr. Rohini Muthuswami’s laboratory, SLS, JNU, for providing access to the fluorescence reader. The project was funded from the SERB-STAR Award Grant (grant no. STR/2021/000045, dated 23/12/2021) from the Government of India. Avinash Kumar Gautam acknowledges the UGC for the Non-NET fellowship and financial assistance from the DBT grant (BT/PR45101/DRUG/134/121/2022). Preeti Umarao thanks the ICMR for the SRF (45/14/2022-BIO/BMS) received during this work.

## CONFLICT OF INTEREST

The authors declare no conflicts of interest.

## DATA AVAILABILITY STATEMENTS

All models of Rho GTPase and their molecular dynamics simulations of all complexes are available in the Zenodo database: EhFP10-Rho GTPase MD simulation Dataset. Zenodo. https://doi.org/10.5281/zenodo.19655410

## SUPPORTING INFORMATION

The supporting information for figures and tables is provided in a separate PDF.

## Footnotes

1 Some EhRho GTPase protein have been referred in literature with alternative names as well such as EhRho2 as EhRacG, EhRho4 as EhRacA, EhRho5 as EhRacD1, EhRho6 as EhRacA2, and EhRho13 as EhRacM. The sequences are compared and there is no difference

## Notes

https://doi.org/10.5281/zenodo.19655410

### Competing Interest Statement

The authors have declared no competing interest.

### Summary of Updates

This version of the manuscript has been revised to correct the spelling of an author's name in both the manuscript file and the submission system metadata.

