## supplementary figures for "Specificity Profiling of the RhoGEF Domain of FP10 with Rho GTPases Involved in Cytoskeleton Remodeling from *Entamoeba histolytica*": SUPPLEMENTARY Rho.pdf (from another version of this preprint: version 1, 2026-05-12)

### SUPPLEMENTARY INFORMATION

**Table S1: Molecular details of seven Rho GTPases**

| Protein Name | Rho1 | Rho2 | Rho4 | Rho5 | Rho6 | Rho8 | Rho13 |
|---|---|---|---|---|---|---|---|
| UniProt ID | P31021 | O76321 | Q24814 | Q24817 | C4M8K6 | C4MA80 | C4M9S3 |
| No. of Residues | 199 | 201 | 196 | 200 | 195 | 202 | 199 |
| Molecular weight ( kDa) | 22.3 | 22.2 | 21.7 | 22.3 | 21.5 | 22.7 | 22.4 |
| pI | 5.66 | 5.96 | 7.49 | 5.78 | 7.49 | 6.16 | 6.89 |
| Extinction coefficient | 21930 | 21680 | 20440 | 21930 | 20440 | 24410 | 20315 |
| Instability index | 46.44 | 32.16 | 36.58 | 46.01 | 38.03 | 28.02 | 30.87 |

**Table S2: Specificity of RhoGEFs by Dbl homology domain containing GEF of FP10**

| GTPase | Activity Rank | Glu33 | Arg4 | Leu69 | Asn43 → | Arg68 → | Other Notable Residues | Summary of Changes |
|---|---|---|---|---|---|---|---|---|
| **Rho2** | **1** | ✔ | ✔ | ✔ | Asn43 | Arg68 | — | Complete conservation of key residues |
| **Rho6** | **2** | ✔ | ✔ | ✔ | **Ser41** | Arg68 | — | Ser41 replaces Asn43 |
| **Rho8** | **3** | ✔ | ✔ | ✔ | — | **Gln72** | — | Loss of Ser41; Arg68→Gln72 |
| **Rho4** | **4** | ✘ | ✘ | ✘ | **Ser41** | **Gln72** | — | Loss of Glu33, Arg4, Leu69 |
| **Rho13** | **5** | ✘ | ✘ | ✘ | **Leu43** | **Gln73** | — | Loss of 3 key residues + Arg68→Gln73 |
| **Rho5** | **6** | ✘ | ✘ | ✘ | **Met48** | **Gln73** | **Asp38** | Complete loss + new substitutions |
| **Rho1** | **7** | ✘ | ✘ | ✘ | **Met47** | **Gln72** | — | Most substitutions and losses |

**Figure S1: Purification of seven Rho GTPase proteins.** 12% SDS-PAGE gel images showing the purified Rho GTPase proteins. The distinct protein bands correspond to the expected molecular weights, as confirmed by comparison with the protein marker ladder.

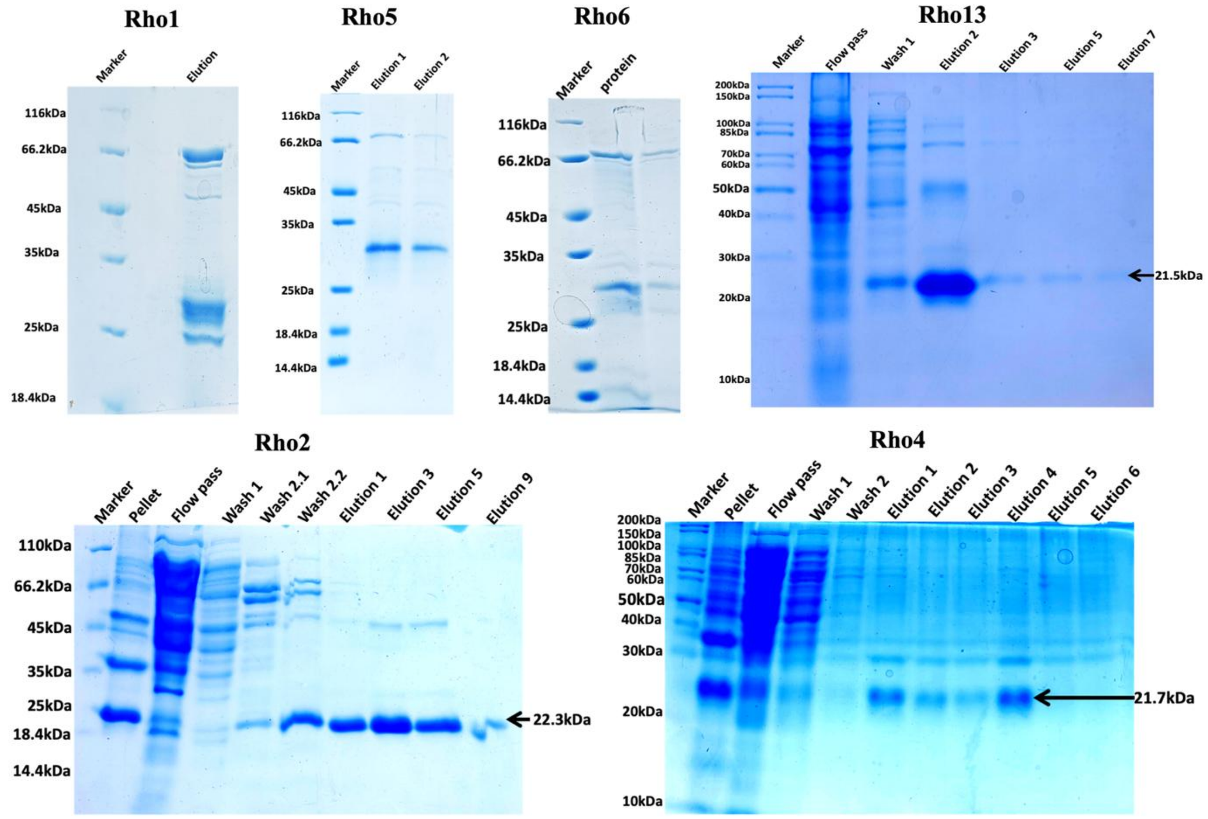

**Figure S2. Structural models of *Entamoeba histolytica* Rho GTPase (EhRho GTPase).** A 3D structure model of six EhRho GTPase is shown in ribbon representation in one orientation generated from AlphaFold3.0. The protein is coloured in a rainbow gradient from the N-terminus (blue) to the C-terminus (red) to highlight overall folding and domain organization. The putative nucleotide-binding pocket is indicated by the bound ligand, corresponding to the conserved GTP/GDP-binding region typical of Rho family proteins. Structural variations among the models primarily reflect differences in loop regions and flexible terminal segments, while the core GTPase domain remains highly conserved. These models are consistent with known structural features of Rho GTPases, including the central β-sheet flanked by α-helices and the switch regions involved in nucleotide-dependent conformational changes.

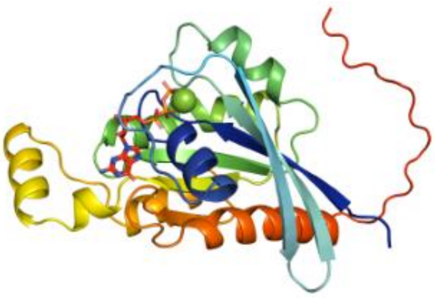

**Rho2_GTP**

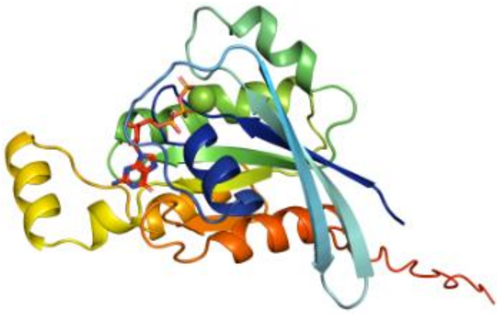

**Rho4_GTP**

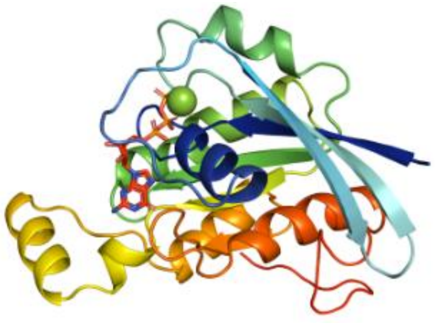

**Rho6_GTP**

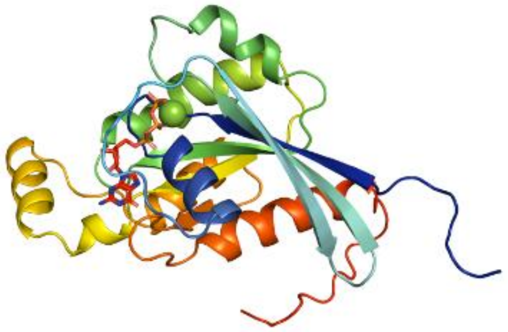

**Rho5_GTP**

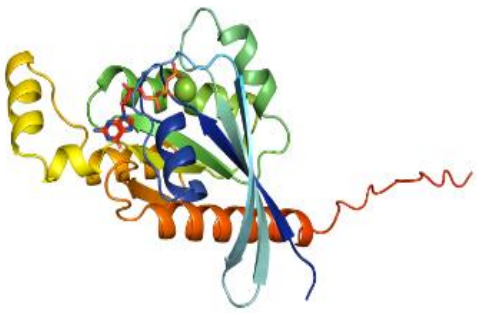

**Rho8_GTP**

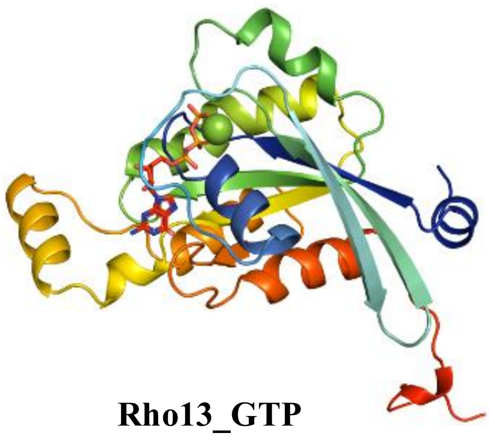

**Rho13_GTP**

**Figure S3. Crystal structure of EhRho1 in the nucleotide-bound state.** Ribbon representations of the crystal structure (GDP-bound form (PDB:3REF) and GTP analog-bound form (PDB:3REG)) of *Entamoeba histolytica* Rho1 (EhRho1) are shown. The protein is colored in a rainbow gradient from the N-terminus (blue) to the C-terminus (red), highlighting the overall fold of the conserved small GTPase domain. The bound nucleotide (GTP/GDP analog) is depicted as a stick representation, and the coordinating $Mg^{2+}$ ion is shown as a sphere within the nucleotide-binding pocket.

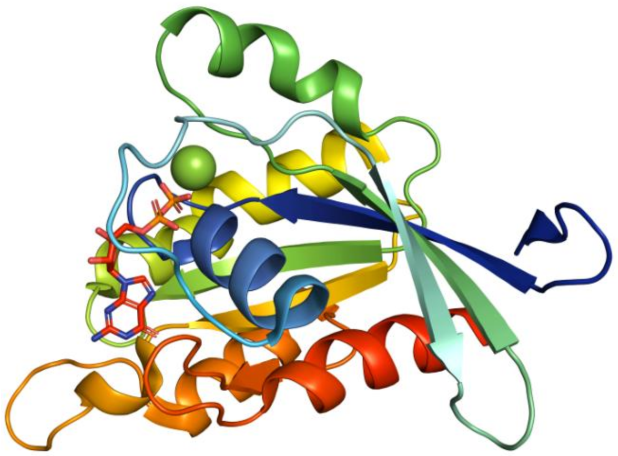

**Rho1_GDP**

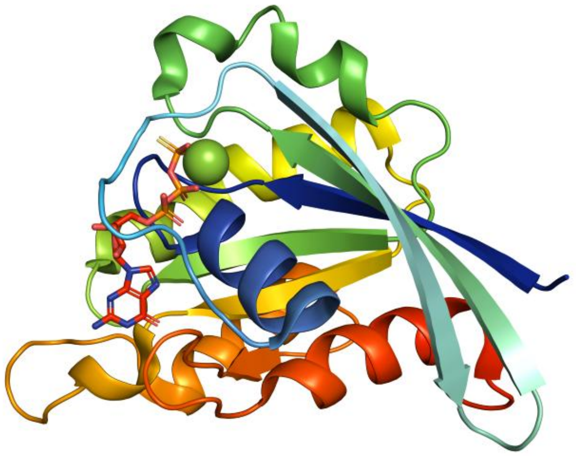

**Rho1_GSP**

**Figure S4: 2D representation of interaction residues in Rho–GEF complexes analyzed via LigPlot. (A)** Comparative analysis of the Rho2–GEF complex with Rho6–GEF and Rho8–GEF reveals minimal differences in the interacting residues. **(B)** In contrast, comparisons of Rho2–GEF with Rho5–GEF, Rho4–GEF, Rho13–GEF, and Rho1–GEF show significant variations in the interaction interfaces, indicating differential binding patterns among these complexes.

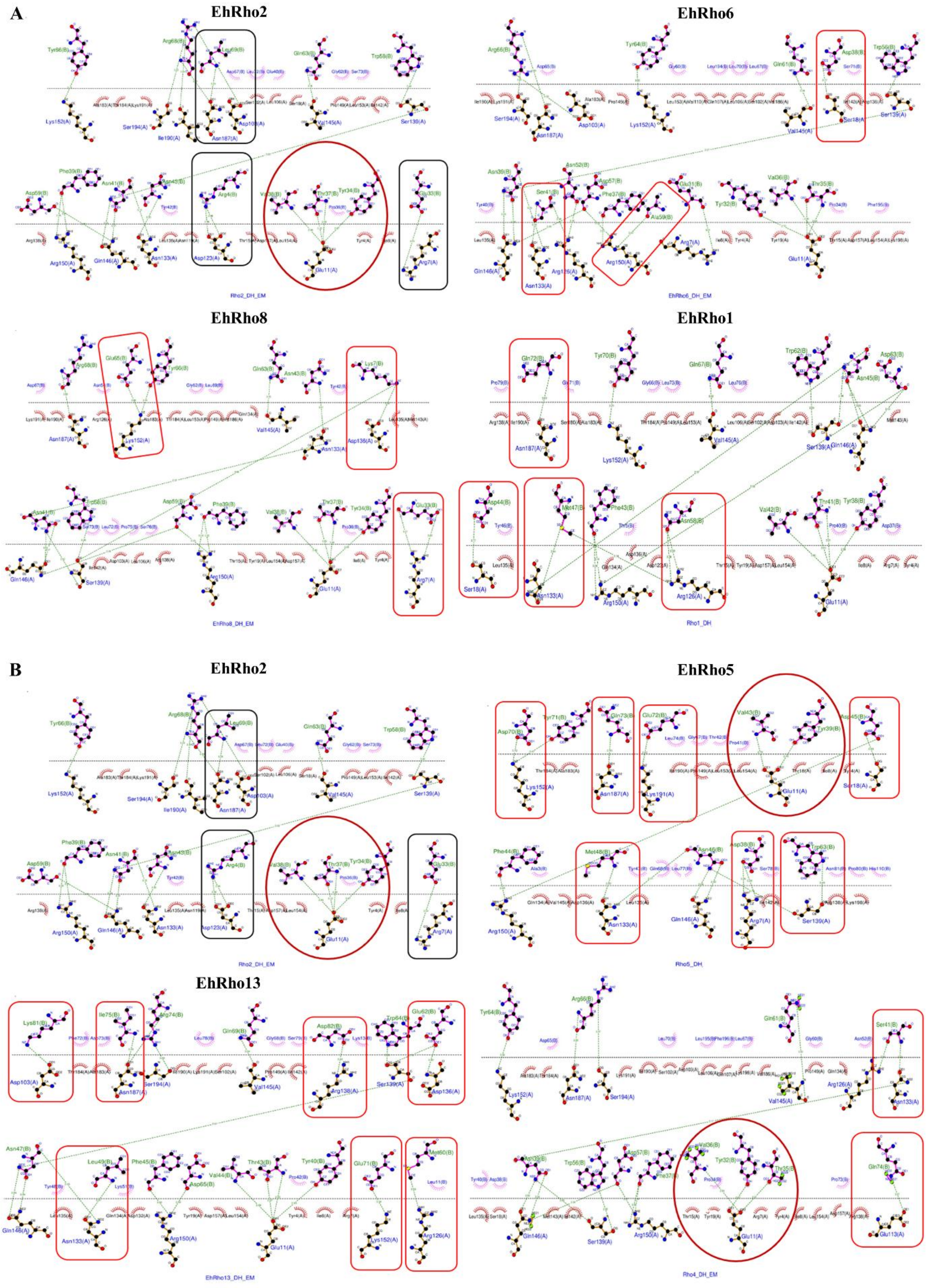

**Figure S5: Analysis of Rho–GEF complexes using the iMODS web server.** Correlation maps showing residue-residue dynamic cross-correlations during complex formation. The maps use a colour gradient in which red indicates a positive correlation, blue indicates a negative correlation, and white indicates no correlation. Rho2–FP10_DH/PH, Rho6–FP10_DH/PH, and Rho8– FP10_DH/PH complexes show dominant red regions, reflecting strong positive correlations between residues. In contrast, the Rho4–FP10_DH/PH, Rho5–FP10_DH/PH, Rho13–FP10_DH/PH, and Rho1–FP10_DH/PH complexes show reduced red colouration, indicating weaker or less correlated residue motions. The overall red intensity follows a gradient from high in the Rho2–FP10_DH/PH complex to low in the Rho1–FP10_DH/PH complex.

**(B)**

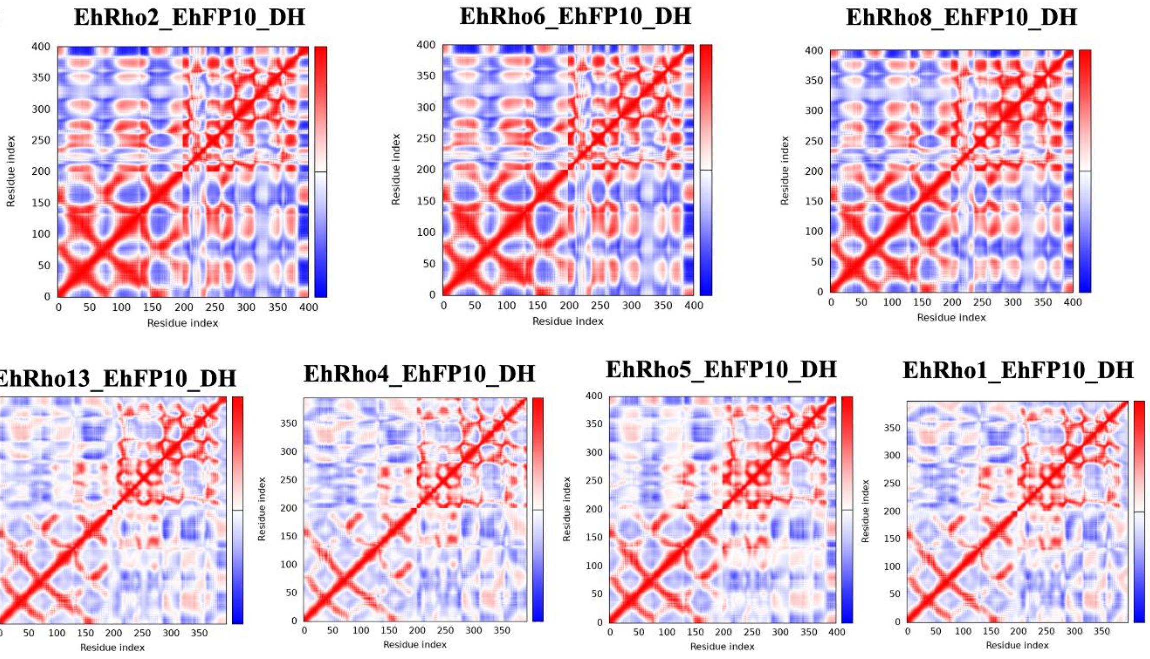
